# Pointing Errors in Non-Metric Virtual Environments

**DOI:** 10.1101/273532

**Authors:** Alexander Muryy, Andrew Glennerster

**Affiliations:** School of Psychology and Clinical Language Sciences, University of Reading, Reading, RG6 6AL, UK

**Keywords:** Human Navigation, Spatial Representation, Virtual Reality, Metric model, Motion Parallax, Binocular Disparity, Topological Model, Labelled Graph, View-Based

## Abstract

There have been suggestions that human navigation may depend on representations that have no metric, Euclidean interpretation but that hypothesis remains contentious. An alternative is that observers build a consistent 3D representation of space. Using immersive virtual reality, we measured the ability of observers to point to targets in mazes that had zero, one or three ‘wormholes’ – regions where the maze changed in configuration (invisibly). In one model, we allowed the configuration of the maze to vary to best explain the pointing data; in a second model we also allowed the local reference frame to be rotated through 90, 180 or 270 degrees. The latter model outperformed the former in the wormhole conditions, inconsistent with a Euclidean cognitive map.

## 1 Introduction

During active exploration of a 3D scene, an observer must build up some kind of representation about the layout of objects that will be useful from a different vantage point. One hypothesis is that the representation corresponds to a type of map with metric measures such as distances and angles conforming to a Euclidean geometry, even if these are not a faithful reproduction of the environment [1–3]. Such a metric cognitive map could be constructed, for instance, on the basis of path integration. It could provide a comprehensive description of the environment and underpin a wide variety of spatial tasks such as general navigation, finding shortcuts or detours and pointing to targets. Crucially, the structure of the data stored in such a map would be independent of the task. However, while there is a predominant view that metric representations may provide an adequate description of small open environments (vista spaces), there is good evidence that this hypothesis does not hold for a global representation of large complex environments [4]. Multiple experimental studies speak against global metric representations [4–9]. For instance, Warren et al [5] found that perceived locations of targets in a labyrinth may overlap, i.e. the perceived geometry of space is not metrically consistent (they report discontinuities in spatial representation, such as “rips” and “folds”). Typically, human participants can navigate efficiently through complicated experimental environments which supports the notion of a global spatial representation. However, their perception of metric qualities is often distorted. For instance, the perceived length of a route depends on the number of turns and junctions it contains [9, 10]. Angular and directional judgments are highly unreliable [7, 11–13] and perceived angles between junctions are biased towards 90° [9, 14].

An alternative type of representation of the environment is a topological one [5, 6, 8, 9, 15–18]. A topological graph consists of a network of nodes and edges that connect them. The exact shape of the path between nodes is not defined because the edges simply represent the connectivity of the environment, e.g. the existence of the path between two locations. Nevertheless, topological knowledge of an environment is sufficient for general navigation and allows the observer to find alternative routes and detours. This type of representation has been used to account for both human and insect navigational behaviour and has been applied in robot navigation systems [16, 19]. Arguments against topological representation for navigation have been raised [20, 21] but, at least in these cases, not convincingly so [22, 23].

Hybrid models, that include both topological information of space connectivity and metric information have also been proposed [5, 8, 15, 24, 25]. Metric information about scene layout is often assumed to be available in the region that is visible from one location, e.g. within a room, which Montello has described these as ‘vista spaces’ [26]. These representations are hierarchical in nature, such that metric representation is reliable locally while at the same time the global metric representation may be distorted to affect both the location of objects and the orientation of local ‘vista spaces’. Evidence compatible with representations of this type has been reported on the basis of pointing, walking or other orientation judgements in environments that were known to the participant [4, 5, 9]. Another aspect of spatial behaviour that is not predicted by a metric map is that the retrieval of information may depend on the observer’s location [14], so that perception of the spatial layout of the scene is different when judged from point A and point B.

In our experiments, we generated virtual environments that were impossible to recreate physically and tested participants’ ability to point to targets that they had encountered previously but could not currently see. The reason that such environments are informative is that they allow predictions to be tested that could not be distinguished in a normal environment. Non-Euclidean environments of this sort were used previously to test human cognitive maps, for example [5, 27–29]. Warren et al [5] created a virtual labyrinth with “wormholes” that teleported participants smoothly between locations. Vasylevska and Kaufmann [28] tested environments with spatially overlapping regions much like the environments in our experiments in order to simulate space compression for VR applications. Zetzsche [29] and Klus [27] developed impossible virtual environments that violate Euclidean metrics and planar topology. Surprisingly, in all these studies human participants showed remarkable insensitivity to metric inconsistencies of space. Warren [5] used ‘as-the-crow-flies’ walking as a measure of the perceived distance and direction of a previously-seen target. They concluded that participants in this task were using a distorted type of map which they called a ‘labelled graph’. The goal of our experiment was similar but, in our case, we compared explicitly a wide range of metric configurations, with and without non-metric variations in local orientation, to see which type of representation could best explain the pointing data.

## 2 Methods

### 2.1 Participants

The 8 participants (3 male and 5 female) were students or members of the School of Psychology and Clinical Language Sciences. All participants had normal or corrected to normal vision, one participant wore glasses during experiment, and all had good stereo-acuity (TNO stereo test, 60 arcsec or better). All participants were naïve to the purpose of the study. Participants were given a one hour practice session in VR to familiarize them with our set-up in a simplified virtual environment (open room with targets in boxes, but no inner walls) and metric versions of the mazes. 6 potential participants (in addition to the 8 who took part) either experienced motion sickness during the practice session or preferred not to continue at this stage. Altogether there were 7 sessions (including the practice) roughly 1 hour each, conducted on different days. Participants were advised not to stay in VR longer than 10 minutes between breaks. They received a reward of 12 pounds per hour. The study received approval of the Research Ethics Committee of the University of Reading.

### 2.2 Experimental set-up

The experiment took place in a 3 by 3m region of the laboratory equipped with Vicon tracking system (14 infrared cameras) that provides 6 d.o.f. information about the headset position and orientation with nominal accuracy ±0.1 mm and 0.15° respectively. We used nVis SX111 head mounted display with 111° field of view and binocular overlap of 50°. A video cable connected the HMD to a video control unit on the ceiling. The position and orientation of the HMD was tracked at 240Hz and passed to the graphics PC with a GTX 1080 video card. The stimuli were designed using Unity 3D system and rendered online at 60fps. Participants were allowed to walk freely and explore the virtual environment in a natural way. The virtual labyrinth was originally a 5 by 5m environment with corridors in the maze 1m wide. In order to fit in the 3 by 3m lab space, the labyrinth was shrunk to 0.6 scale (e.g. 60cm wide corridors) which meant that the floor was displayed about 1m below eye height. Participants generally found this acceptable and did not notice that the room was not normal size, compatible with previous experiments [30]. During the experiment, participants wore a virtual wristband that provided information about the task (shown, for illustrative purposes only, at the top of Fig. 1b). Participants used a hand-held 3D tracked pointing device to point at targets during the pointing phase. In VR, the pointing device looked like as a small sphere (R=5cm) with an infinitely long ray emanating from it in both directions; the text was displayed next to the hand in VR providing instructions (e.g. ‘point to Red’).

**Fig. 1.**
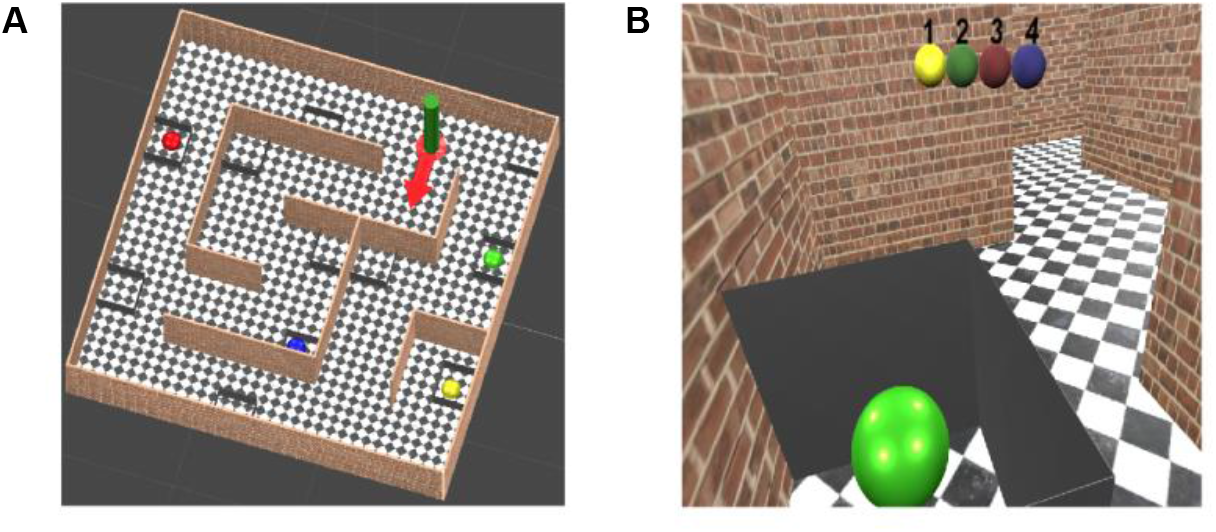
Views of the labyrinth. A) View from above. B) First person view. The green target is visible inside a grey box. The target sequence is shown on top (Y-G-R-B) and the current target is highlighted (Yellow).

### 2.3 Stimuli

We designed 2 general layouts of the virtual labyrinth. Scene 1 is shown in Fig 1 and 2. The virtual environment could be subdivided into 25 (5×5) elementary squares each having a size equal to the corridor’s width. A green cylinder (Fig. 1a) indicated the starting location. In order to start the experiment, the participant should stand at the start location and look along the corridor’s width. In order to start the experiment, the participant stood at the start location and looked along the direction indicated by the red arrow. Then green cylinder and red arrow disappeared, so that the starting location was not marked during the exploration phase. The labyrinth contained 4 target objects (red, green, blue and yellow spheres) hidden inside open grey boxes, so that they could be seen only from a short distance. Other empty grey boxes were added as distractors. Fig. 1b shows a first-person view along a virtual corridor containing the green target inside a grey box. Participants wore a virtual wristband that displayed information about the current task, namely the sequence in which targets should be collected (e.g. Yellow-Green-Red-Blue, see Fig. 1b, top) and then pointed to. See Supplementary Information for movies of the experiment and plan views of the scenes.

**Fig. 2.**
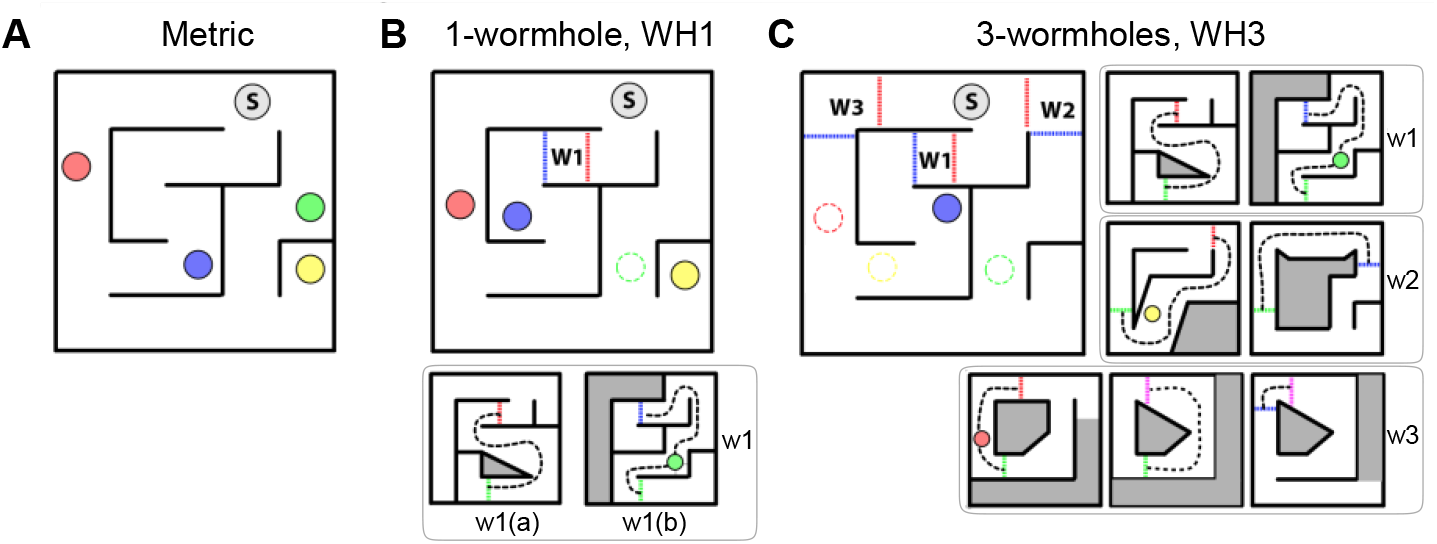
Schematics of the labyrinth. A) Metric condition. B) Non-metric 1-wormhole condition. The green target was placed inside the wormhole. C) Non-metric 3-wormholes condition. The general layout (containing Start, which is marked as ‘S’) remained constant between conditions. The wormholes are marked with letters W surrounded by red and blue triggers. As the participant crossed a trigger, the environment changed as shown in the sub-schematics. Inside a wormhole, the participant could only walk along the route marked by black dashed line. These are for Scene 1. Equivalent set of maps were generated for Scene 2 (see layout in Fig. 10 and in Supplementary Information).

For each labyrinth, we increased the complexity of the environment by extending the length of the corridors with non-metric ‘wormholes’, see Fig. 2b and c. There were 3 conditions per scene: one metric, one containing 1-wormhole and one containing 3. Start is marked as ‘S’, colored circles indicate targets. The dashed lines acted as invisible triggers: when a participant crossed the trigger the environment changed as shown in the sub-plots. For instance, in the 1-wormhole condition shown in Fig. 2b, when a participant crossed the red trigger the environment changed to schematic W1(a); as the participant continued walking down the path through the wormhole (the accessible region inside a wormhole is marked by the dashed black line) and crosses the green trigger, the environment changed again to schematic W1(b), then as the participant crossed the blue trigger he or she would exit the wormhole and the environment change back to the general layout. Participants were not aware of the presence of the triggers because the local views of the environment did not change during the trigger crossing.

For Scene 1, the same original or ‘base level’ layout applied in all three conditions (metric, 1-wormhole and 3-wormholes), as shown in Fig. 2a. Similar principles applied to Scene 2 (which also had a metric, one-wormhole and three-wormhole version). The corridors through the wormholes did not have junctions, thus the topological connectivity of space was the same in all 3 conditions. The only difference between metric and non-metric conditions is the length and configuration of the corridors. The wormholes extended the corridors in a way that made a global metric representation impossible. For instance, the path through the wormhole in Fig. 2b has the shape of the figure of eight, i.e. it crosses itself, although there are no visible junctions along that path, which is physically impossible.

### 2.4 Procedure

Participants were asked to perform a navigational and pointing tasks in a virtual labyrinth (see Fig. 1) although we report here only on the results of the pointing task. A single experimental session that took about 1 hour during the three conditions were tested sequentially metric, 1-wormhole and 3-wormholes, all with the same general layout (i.e. all Scene 1 or Scene 2). This helped participants to cope with the more complex environments. The tasks and instructions were identical for all 3 conditions. For each condition, participants had to complete 8 rounds of walking and pointing. Each round started with a navigational task during which the participants were asked to collect all 4 target objects in a specified order in the most efficient way. The meaning of ‘efficient’ was left to participants to decide: it could mean choosing the shortest path, or the smallest number of turns or junctions (i.e. navigational decisions) but they were told not to hurry. After all targets were collected, the participants were instructed to stay at the location of the last target and to point to the Start location and then all other target locations twice, in a specified order (Start-R-G-B-Y and repeat), using a pointing device. Apart from the last sphere, the targets could not be seen from the pointing location, hence the participants had to rely on their spatial representation of the environment to complete the pointing task. Once the pointing task was complete, the next round of walking and pointing commenced. During all 8 rounds the target locations remained constant. The first 5 rounds were a ‘learning’ phase in which participants always began at the Start location and collected targets in the same sequence Start-Red-Green-Blue-Yellow (S-R-G-B-Y) and finally pointed to all the targets from Yellow (i.e. from the last target). After pointing, the walls of the labyrinth disappeared and the participant went directly to the Start location and begin the next round. The purpose of the learning phase was to allow participants to build up a spatial representation of the labyrinth gradually through multiple repetitions of the same navigational task. During the test phase (the last 3 rounds) the navigational sequences were changed to three new sequences: Y-G-B-Y-R, R-B-R-Y-G and G-Y-G-R-B, so that during test the participants had to solve novel navigational tasks. They did not have to go to the Start locations at the beginning of a round but instead started at the point where the previous round ended. Importantly, the pointing locations at the ends of the sequences of the test rounds included all target locations except for Yellow, which was the pointing location during the learning phase. Thus, during test phase we collected pointing data from all target locations, except for Yellow.

Excluding the practice session, each participant carried out 6 experimental sessions, each on a different day. We tested one Scene per session (Scene 1 or Scene 2) with three repetitions of each Scene. Across repetitions, the structure of the labyrinth and target locations were identical, but the colours of the targets were changed. This meant that while the instructions remained the same (e.g., in the learning phase, collect targets in sequence R-G-B-Y) the actual paths relating to those tasks were different on different repetitions. In the end, in Scene 1 we had 3 sets of pointing data in the test phase from the location marked as red in Fig. 2 and 2 sets of pointing data from green, blue and yellow locations (in scene 2: 3 sets from blue and 2 sets from the others). Altogether this constituted 72 pointing vectors per scene per condition per participant: 24 vectors from the red target and 16 vectors from green, blue and yellow.

## 3 Results

During the first couple of learning trials, while participants were not familiar with the structure of the labyrinth, their paths appeared relatively random, but closer to the end of the learning phase the participants could navigate more systematically. Here, we present the results of pointing in the test phase, after participants had completed the 5 learning trials. Fig. 3 illustrates the paths that participants took through the maze during the test phase, after the learning phase of 5 rounds. In the cases shown in Fig. 3, participants were always navigating from the blue sphere to the yellow sphere, for a ‘metric’ (normal) maze, a maze with one wormhole and a maze with three wormholes. There are 4 possible solutions to this task, assuming that participants did not double back on themselves and avoided ‘loops’ in their trajectory (returning to a junction) and, in fact, by the test phase participants did not do this, demonstrating learning during the first phase. The shortest route is marked in red (in these cases, the shortest topological route is also the shortest metric route). Notice that in the 3-wormhole condition the participants’ paths are significantly longer and more complex than in the metric and one-wormhole conditions even when, topologically speaking, they take the same route as in the metric condition. Participants often did not notice the non-metric nature of WH1 condition although they sometimes reported that the Green target (inside the wormhole) was harder to find. In the three-wormhole (WH3) condition all participants were aware that the scene was geometrically impossible.

**Fig. 3.**
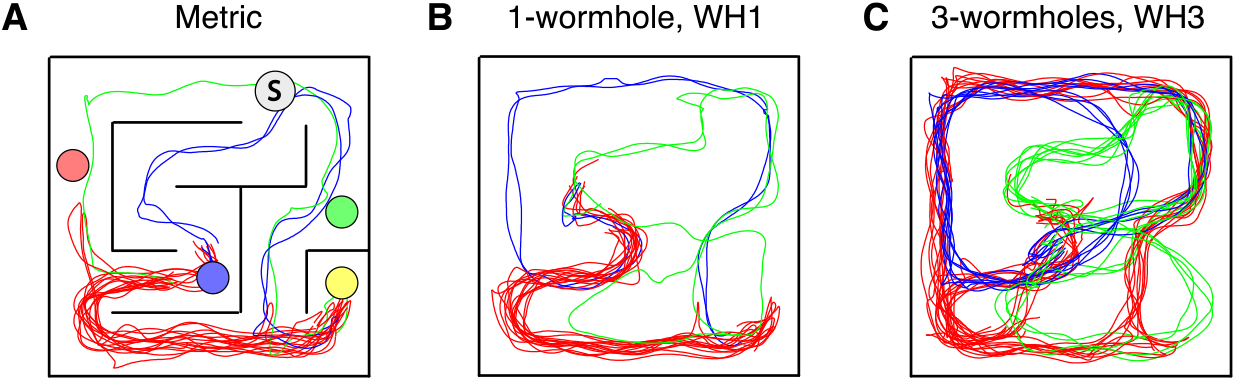
Example of a participants’ paths during one subset of the task, “go from B to Y”. The full task for this round was: start at Yellow and go to targets in the order G-**B-Y**-R. The shortest path is marked in red, while green and blue lines represent alternative routes. Notice that in the 3-wormhole condition paths are significantly longer.

Fig. 4 shows pointing data for the test phase in Scene 1 (8 participants, 3 conditions and 3 repetitions). The plot shows signed pointing errors, where positive errors are counter-clockwise relative to the true direction of the target from the centre of the hand-held pointing device. Because of the order in which the pointing zones were visited, the number of ‘shots’ from each pointing zone was not identical: in scene 1 there were 48 pointing data-points from the red location to each of the other targets, and 32 data-points from green, blue and yellow locations while in scene 2 (see Supplementary Information): there were 48 pointing data-points from the blue location and 32 data-points from red, green and yellow locations.

**Fig. 4.**
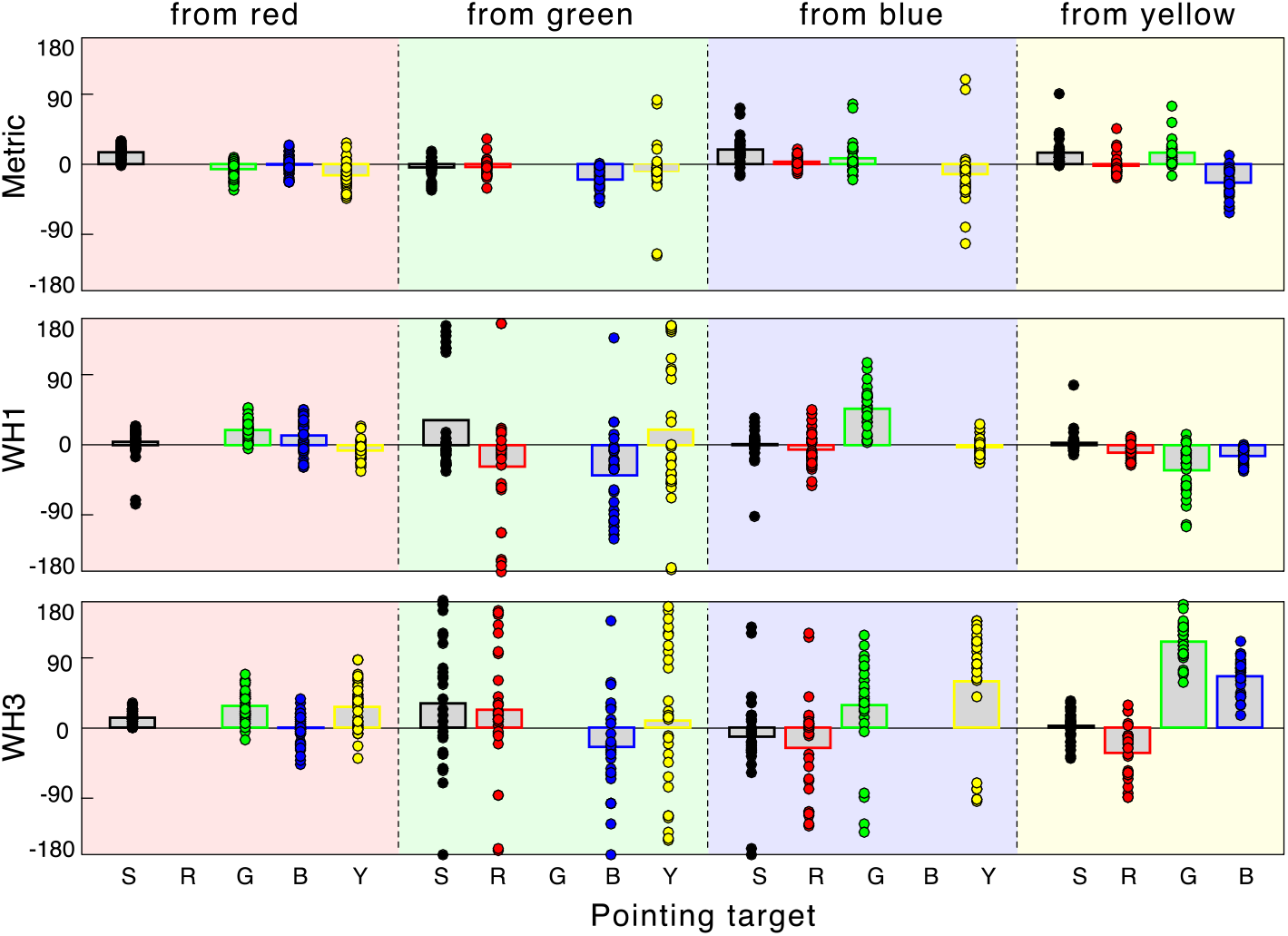
Signed pointing errors for all participants (n=8) and all repetitions (3), for Scene 1. Data for the three conditions (metric, 1-wormhole and 3-wormhole) are shown in the top, middle and bottom rows. Pointing errors are shown individually with coloured symbols indicating the pointing target (Start, Red, Green, Blue and Yellow), the bar height indicating the mean error in each case. The plot background colour indicates the location from which the participant pointed (red, green, blue and yellow columns). Very similar data were obtained for Scene 2.

Fig. 4 clearly shows that the magnitude of pointing errors increases from metric to 1-wormhole to 3-wormhole conditions. This is summarized in Fig. 5a which shows the standard deviation of all the pointing errors (across all participants, repetitions and scenes) for these conditions. However, it would be misleading to describe this as simply an increase in noise or non-specific variability. Fig. 5b illustrates one example of why this is not the case. The solid lines and dotted lines show pointing directions measured on two different days, demonstrating that the directions can be quite repeatable across days. This makes the errors in the wormhole condition all the more remarkable, since the errors are often close to 180 degrees. In this case, it appears that the participant might have been disoriented, but in exactly the same way on both days. However, the pointing vectors cannot be explained by rotation alone – there is a roughly 90° angle between the red and blue pointing vectors while in reality blue and red targets were located on the same line when viewed from green. We discuss some possible ways to model this behaviour in the model section.

**Fig. 5.**
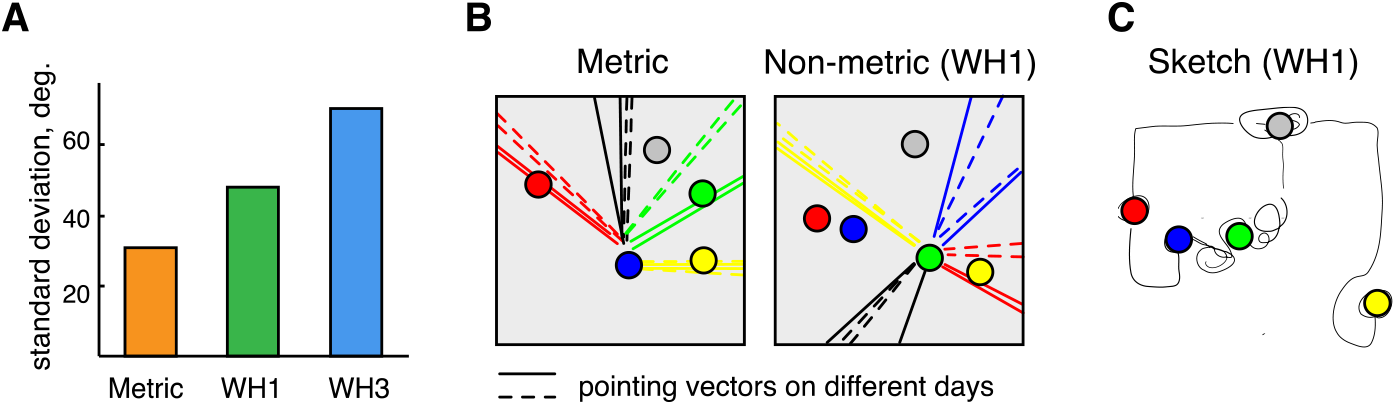
Summary of pointing errors. A) The standard deviations of signed pointing errors across all participants and all conditions for both Scenes. B) Examples of pointing vectors in a metric (left) and non-metric condition (1-wormhole). In the non-metric case, the green target was inside the wormhole. C) A sketch drawn by a participant for the WH1 condition, scene 1. Notice that the participant is clearly much less confident about the path to the Green target, which is inside a wormhole. More sketches are included in the Supplementary Information.

Such distortions are not limited to a single participant. Fig. 6 shows pointing vectors of all participants to the green target from all pointing locations (left) and the same for the yellow target (right) in a one-wormhole condition (illustration from Scene 2, see Fig. 10 and Supplementary Information for schematics). Just like in Fig 5b, the green target was inside a wormhole, while the red, blue and yellow targets and the Start location were not, and hence it is no surprise that the pointing to and from the green target shows particularly large errors. For example, pointing from the green to the yellow target and from the yellow to the green target both had errors of about 180°. Despite these large errors, participants point accurately to the yellow target from both the red and blue locations which is understandable as neither are in a wormhole and they are connected by ‘metric’ (physically possible) routes.

**Fig. 6.**
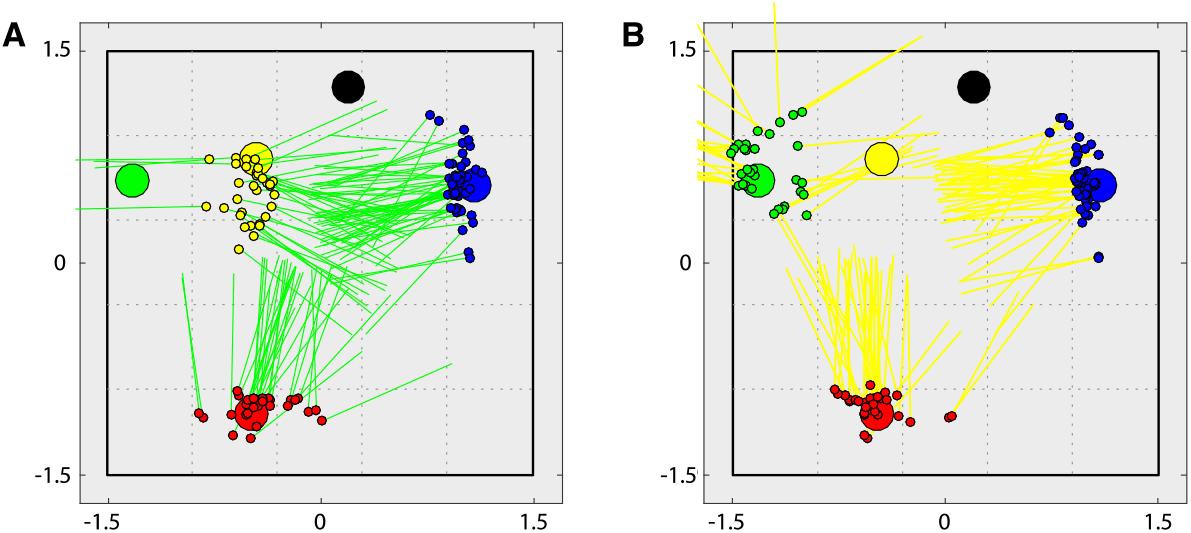
Example of systematic pointing errors in a non-metric condition with one wormhole. A) Pointing vectors to the green target from each of the other locations. The small circles show the hand location. The green target is inside the wormhole (see schematic in Fig. 10a). Most participants wrongly report that the green target is at the centre of the maze. B) Pointing vectors to the yellow target; these are reasonably accurate from the red and blue locations but there are systematic errors from the green location. These data are from Scene 2.

## 4 Model

The systematic pointing errors that participants make in non-metric environments indicate that they are not basing their responses on a correct global spatial representation of the target locations. Note that ‘correct’ locations of each target can still be defined according to path integration, even in the wormhole conditions. Two qualitatively different kinds of distortions are possible: structural distortion and/or local orientational distortion where the orientation of local reference frames can be misestimated (Fig. 7). In order to separate these hypotheses, we estimated the optimal configuration of targets that could best explain the pointing data in the sense that it maximizes the following likelihood function:

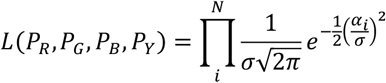

where *P_R_, P_G_, P_B_* and *P_Y_* are locations of the targets and *α_i_* is the pointing error with respect to the actual direction of the targets; the hand location for any given trial was rigidly translated with the pointing location. This function returns the largest value when angular errors, *α_i_*, are zero.

**Fig. 7.**
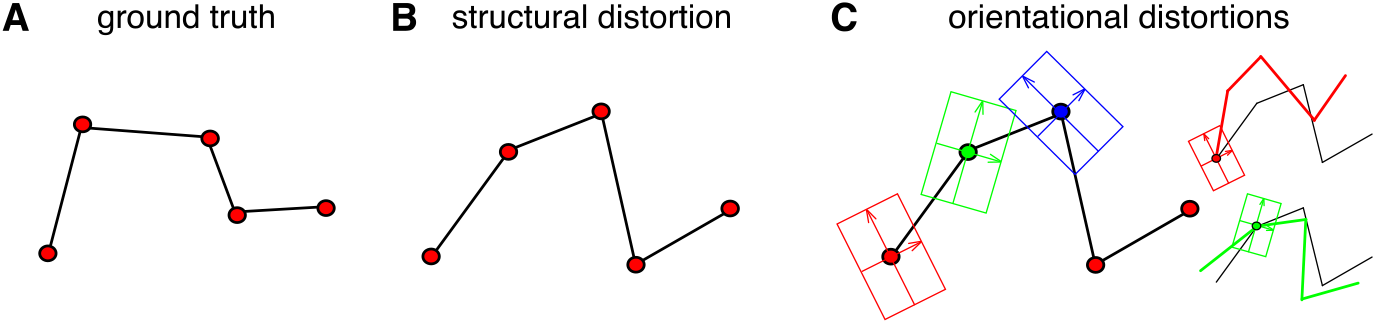
Possible distortions of spatial representations. A) Ground truth structure. B) A structural distortion compared to A). C) Here, in addition, there is orientational distortion so that local reference frames are oriented inconsistently. The insets on the right show the consequences of the local reference frame having a changing orientation, e.g. on pointing responses.

Altogether, there were *N* = 144 pointing vectors (48 from Blue, and 32 from Red, Green and Yellow) per participant per condition. We set the angular standard deviation, *σ*, as 50° but repeating the analysis with different values showed that the locations returned were rather insensitive to the value of *σ*. The Start location was treated as fixed in the optimization (it was, incidentally, always at a fixed location in the laboratory). The set of potential target locations was discrete and limited to the centers of a 5×5 grid with the same size as the labyrinth (see Fig. 8). We optimised locations of the targets (parameters *P_R,G,B,Y_*) in order to maximize the likelihood *L*. The optimization was done per condition per participant and data from the 3 repetitions were combined together i.e. we assume that on different repetitions participants develop similar representations.

**Fig. 8.**
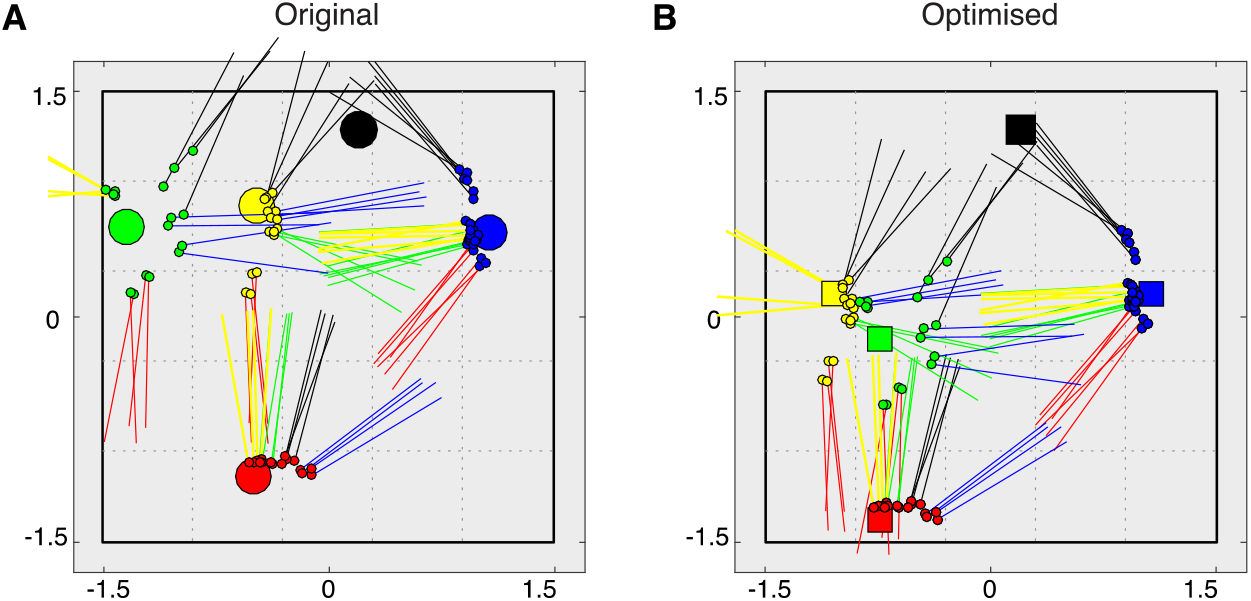
Example of a distorted spatial map explaining pointing data in a 1-wormhole condition. A) The original configuration of targets and the pointing vectors from 3 repetitions for a single participant. Taking ground truth as the ‘model’, the log likelihood, log*(L)*, of these pointing data is −105.3 (see text). B) The optimised (distorted) configuration of targets that best explains the pointing data. Now log*(L)* = −58.2 (see text). Notice that the locations of the green and yellow targets are swapped relative to their locations in (A). These data are for Scene 2.

Fig. 8a shows the original pointing vectors from a single participant in Scene 2. The log-likelihood of the pointing data for this configuration according to our model is log(L)=−105 whereas, when the configuration of the targets is optimised this improves to log(L) =−58 (see Fig 8b). Notice that the green and yellow targets have now (roughly) swapped places, consistent with the pointing directions shown in Fig 6. Finally, in order to capture the orientational distortions illustrated in Fig 5b, we additionally allowed a rigid rotation of the pointing vectors that a participant made from each pointing location. These rotational parameters, *ρ_r_, ρ_g_, ρ_b_, ρ_y_*, describe the orientations of the local reference frames around the red, green, blue and yellow pointing locations respectively and could only take discrete values of 0°, 90°, 180° and 270° because these were the only directions that a participant could face while looking along a corridor. The rotational parameters were optimised at the same time as the positional ones. Separate optimisations were carried out per participant (n=8), per scene (n=2) and per condition (metric, one-wormhole and 3-wormhole). Each of the 4 locations was encoded by 2 parameters (*x, y*) thus, altogether for the first model we had 8*2*4*2= 128 parameters per condition. In the second model, there were 8*2*4*3 = 192 parameters per condition since each location was now encoded by 3 parameters (*x,y,ρ*) rather than just (*x,y*). The total likelihood of all the data under each of the two models was calculated by multiplying individual likelihoods per ‘shot’. Adding more parameters to a model inevitably improves the likelihood of the best fit of that model, so in order to compare models with different numbers of parameters we calculate information criterion:

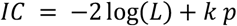

where *L* is the total likelihood of all data, *p* is the number of parameters (0 for original data, 128 for the translation model and 192 for translation and rotational model) and *k*=2 by definition for the Akiake Information criterion (AIC) and *k*=log*(n)* for the Bayesian Information Criterion (BIC). These are shown in Fig. 9a and b for the original (ground truth) configuration and for these two optimisation two models. In the ‘metric’ (normal) maze, assuming the ground truth location of the targets is the best model. However, in the one-wormhole condition the pattern for AIC is reversed and for the three-wormhole condition the reversal is present for both AIC and BIC values. In this case, the best explanation of the data is one that not only allows for a distortion of the location of the targets in the participant’s memory but also assumes that their reference frame for pointing may be rotated independently for different pointing locations.

**Fig. 9.**
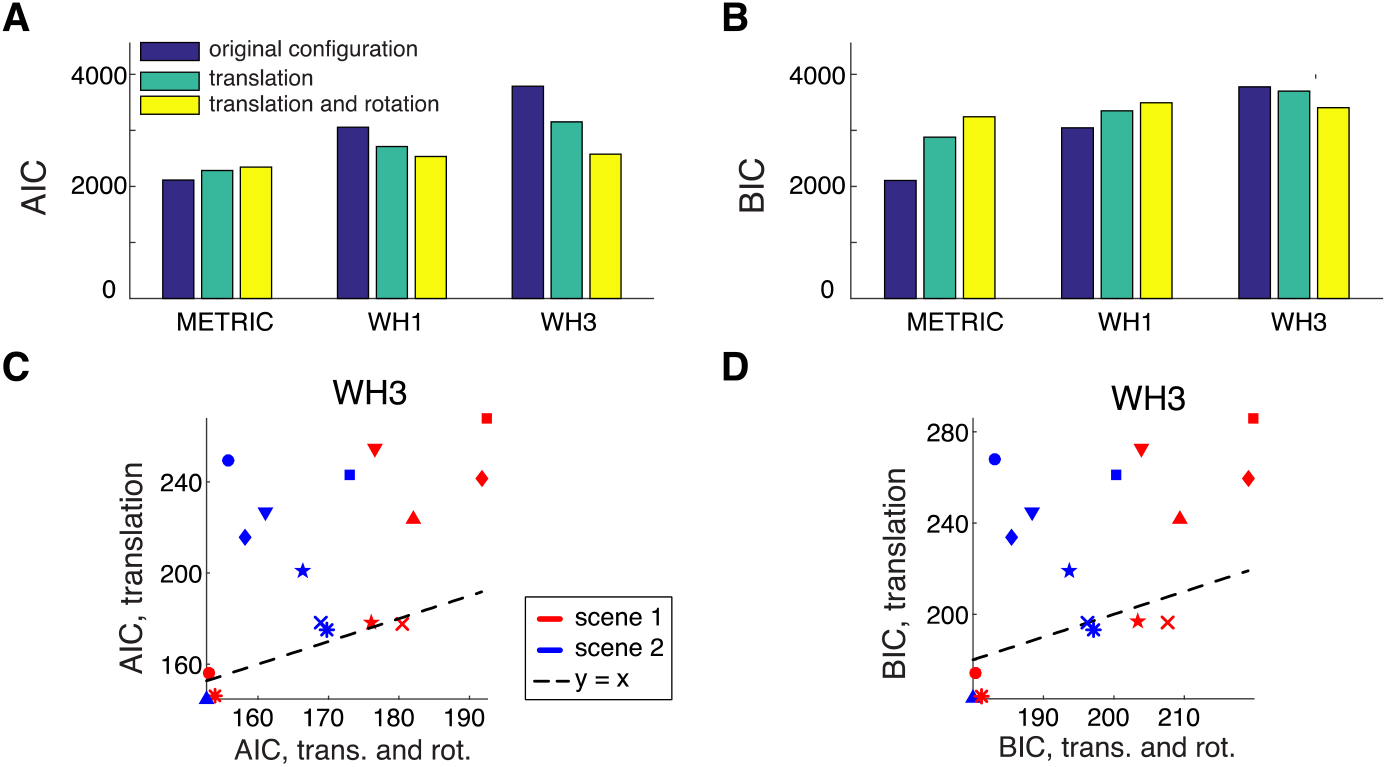
A) Akaike Information Criterion values are shown for each condition under 3 models: ground truth, optimised translation and optimised translation-and-rotation). B) Same for Bayesian Information Criterion. C-D) compare the AIC and BIC values for the translation and translation+rotation models. Different symbols for each participant and colours for each Scene.

## 5 Discussion

Our results show that adding wormholes to a labyrinth environment increases pointing errors (Fig. 5a) in a way that is partly explained by assuming that observers build a distorted model of the environment (Fig. 8) but more fully explained by assuming that observers also rotate their local reference frame when they are at certain locations (Fig. 9). The latter explanation is not compatible with a single globally consistent 3D representation of the environment since that could not tolerate different rotations for different parts of the scene.

The argument that ‘visual space’ is distorted is a familiar one, although much of the literature on this refers to the perception of an observer that is either stationary or only moving a few centimetres [31–34]. It is more unusual to try and find a distorted map of an environment to explain an observer’s behaviour when the observer is free to move around and explore all of that environment, partly because in this situation it is not immediately obvious what systematic distortions could apply. Nevertheless, in the peculiar wormhole environments that we have examined, systematic distortions of the scene layout do provide a better explanation of pointing behaviour than the ground truth layout (Fig. 9).

The modelling we have done is agnostic about the causes of the distortions. One speculation is shown in Fig. 10. This illustrates the same maze layout as is shown in Fig. 6 and Fig. 8 where participants made large errors in pointing from the green to the yellow target and vice versa. The green target is located inside a wormhole next to the left wall and hence is, in reality, to the left of the yellow target. However, as Fig. 8b shows, the best explanation of participants’ pointing is that they believe the green target to be near the centre of the maze (i.e. to the right of yellow in these plan views). As Figure 10 shows, this would be the case if participants’ conception of the wormhole was that it was ‘squashed’ into a region between the entrance and exit of the wormhole in the rest of the maze. Participants might do this because they have learnt that the red, yellow and blue spheres are near the outer walls, forcing an interpretation in which the wormhole region is within the central part of the maze.

**Fig. 10.**
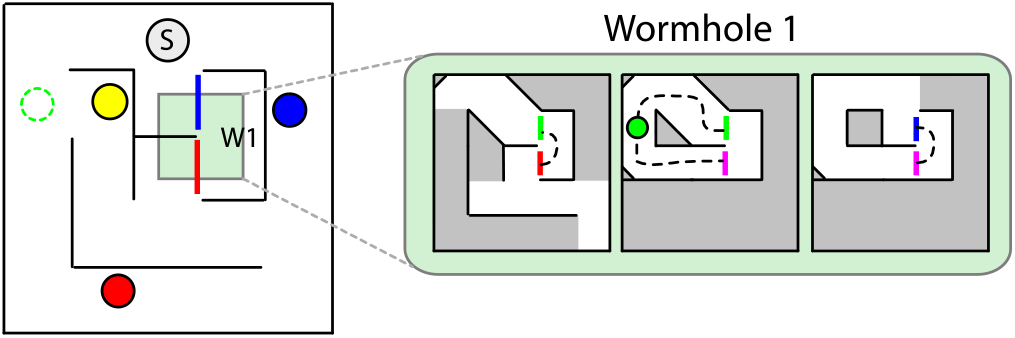
A speculation about a possible cause of the bias shown in Figs 6 and 8. The corridors making up the wormhole are shown in the green panel on the right. These are shown ‘squashed’ into the location where the entrance and exit to the wormhole lie (left). This process leaves the green target to the right of the yellow target in the resulting map. This example is from Scene 2.

Fig. 10 does not include any distortion of local orientation. When we add this in as a variable (Fig. 9), there is no longer a single, consistent metric representation that corresponds to the output of the model (i.e. *x, y* and *ρ* for each target location). This fits with the conclusion of a range of studies based on navigation or spatial knowledge of environments that find evidence against a single consistent representation of 3D space [7–9, 14, 24, 29, 35–37]. For example, Warren et al [5] claimed that there was no consistent representation of space that could account for their data (where the task was to walk directly to targets that had been experienced in a maze with wormholes, i.e. very like our pointing measure but with an indication of perceived distance as well as direction). Our experiment and analysis build on the conclusion of Warren et al (2017) who advocate the idea of a labelled graph. In particular, we make a direct comparison between three models (ground truth, a distorted map or a translation-plus-rotation model) and find that the translation-plus-rotation model fits best. This is compatible with the idea that in the real world, especially in complex environments, observers learn a topological structure first and then a progressively more accurate labelled graph as they become more familiar with the terrain until, eventually, the information about each edge of the graph is so accurate that the result is, at least in theory, impossible to distinguish from a consistent, metric map. This putative hierarchical calibration process for representing space is similar to hypotheses about observers’ representation of surface shape from binocular disparity [38–40].

## Acknowledgements

This research was supported by EPSRC/Dstl grant EP/N019423/1.

## Supplementary Information

Additional figures, movies and raw data are available at: http://www.glennersterlab.com/MuryyGlennerster2018_SupplementaryInfo.zip

